# Generation of an induced pluripotent stem cell line from a healthy adult indigenous Nigerian participant

**DOI:** 10.1101/2023.07.21.550059

**Authors:** Zaid Muhammad, Phoebe W. Brown, Larema Babazau, Abdulrahman I. Alkhamis, Baba W. Goni, Haruna A. Nggada, Kefas M. Mbaya, Selina Wray, Isa H. Marte, Celeste M. Karch, Louise C. Serpell, Mahmoud B. Maina

**Author notes:** Correspondence: Maina M.B. &.

## Abstract

Genetic backgrounds contribute to cellular phenotypes, drug responsiveness, and health outcomes. However, the majority of human induced pluripotent stem cell (iPSC) lines are derived from individuals of European descent. Thus, there is a major, unmet need in the generation, characterisation, and distribution of iPSCs from diverse ancestries. To begin to address this need, we have generated iPSCs from dermal fibroblasts isolated from a healthy 60-year-old indigenous Nigerian male belonging to the Babur ethnic group. The iPSCs were generated using Sendai virus, and copy number variation (CNV) analysis revealed no new major abnormalities compared to the parental fibroblasts. The iPSCs have been characterised for pluripotency markers and morphology and successfully differentiated into neural progenitor cells and astrocytes. This iPSC line could serve as a healthy control in comparative studies and can be used in disease modelling, toxicity assessments, genetic analyses, and drug discovery processes within an African genetic background. To bolster the inclusion of African models in biomedical research, this iPSC line will be made available to the broader scientific community. Ongoing efforts focus on generating more lines from diverse indigenous populations towards creating a dedicated open-access African iPSC biobank.

## Resource Table

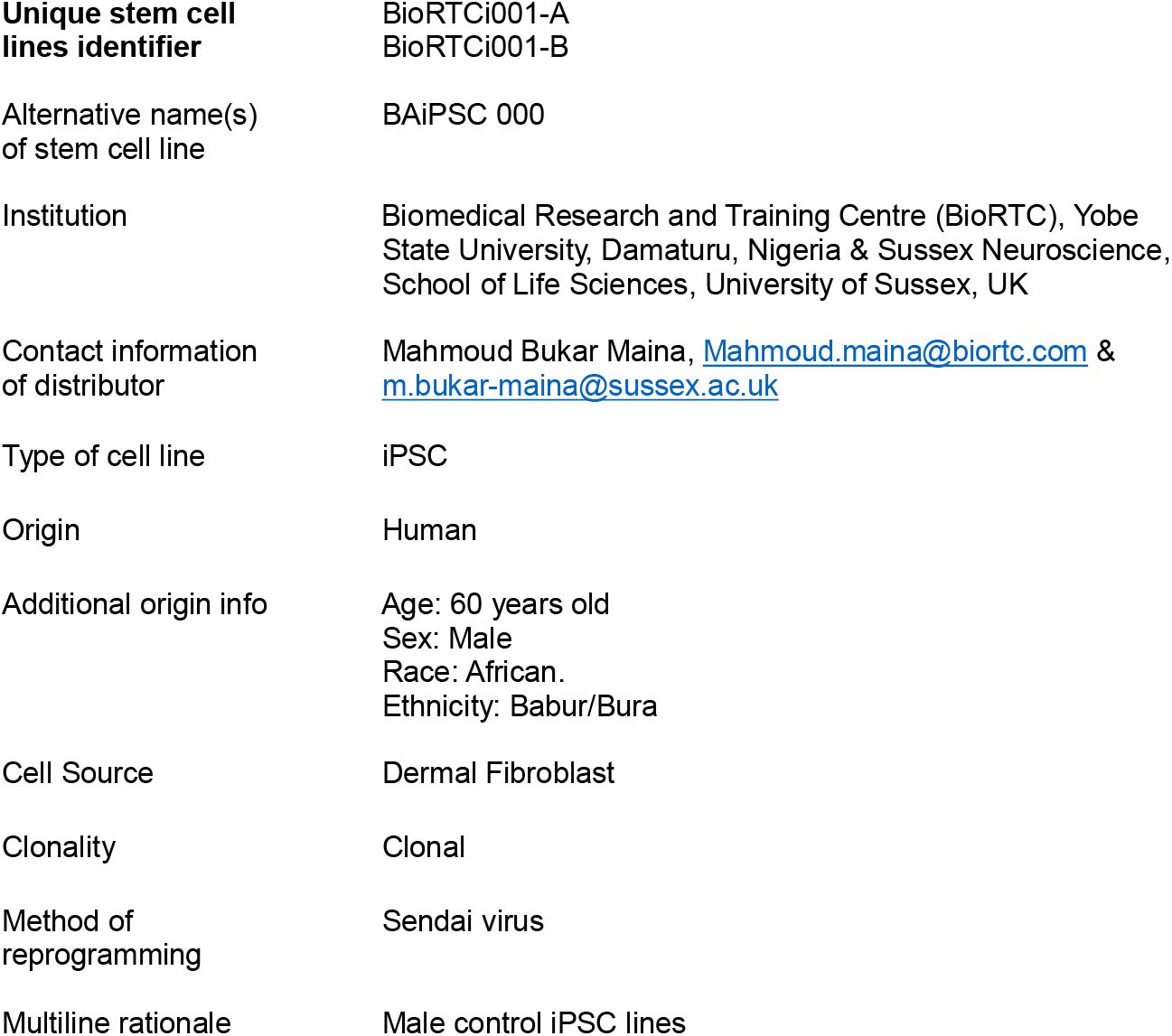
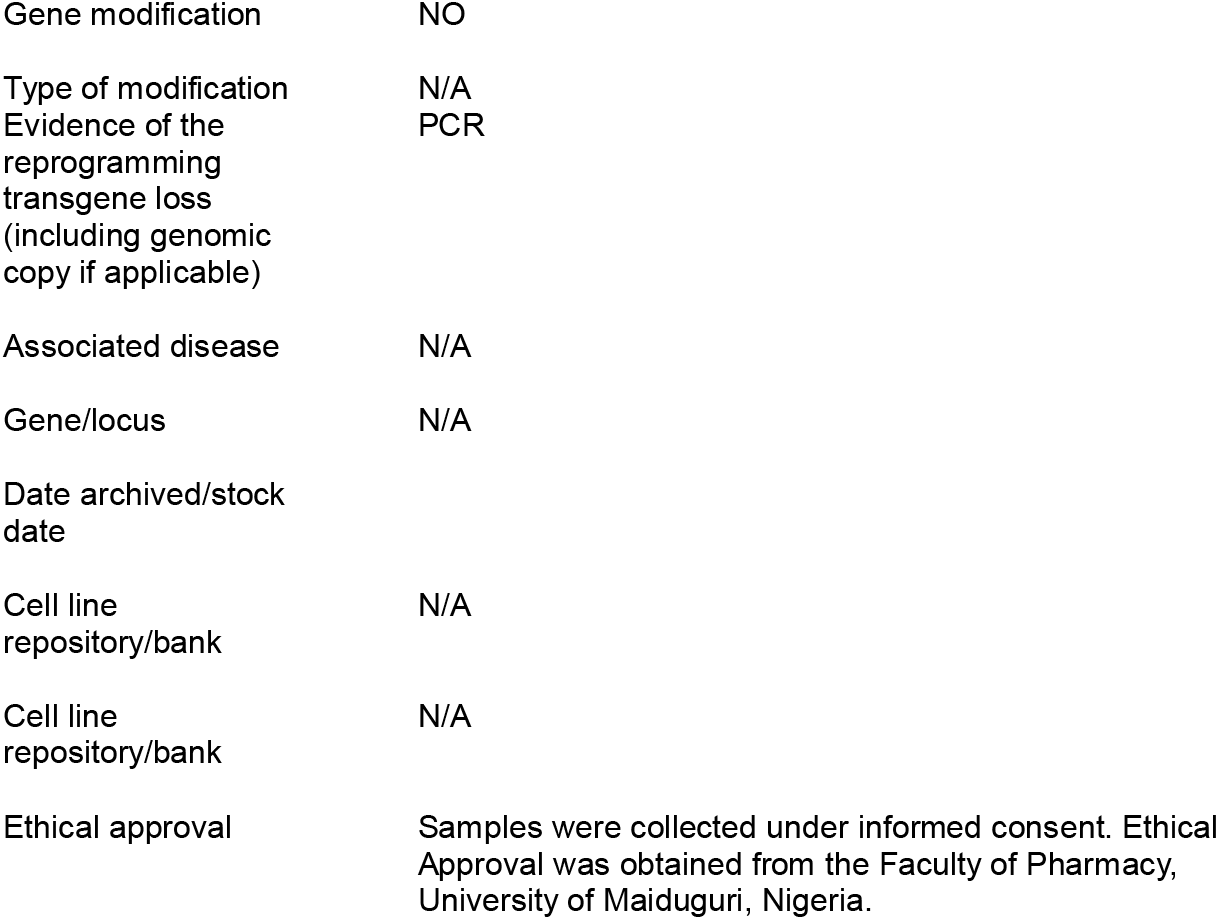

## 1. Resource Utility

The African population harbours the highest level of genetic variation among all populations [1, 2]. These genetic differences impact a range of biological phenomena, from cellular mechanisms to pharmacological responses. For instance, subtle genetic variations can lead to differences in the expression of quantitative trait loci (eQTLs), influencing disease susceptibility and progression [3], including Alzheimer’s disease [4]. Hence, including diverse populations in biomedical research is critical, particularly those with high genetic diversity, like the African population. Our newly derived iPSC line from an indigenous Nigerian represents an invaluable resource for various biomedical investigations, including disease modelling, toxicity assessments, genetic analyses, and drug discovery processes within an African genetic background. This iPSC line can also serve as an essential control for comparative studies across diverse ethnic groups, thus advancing more equitable representation in genetic research. Moreover, this iPSC line can be manipulated with gene editing technologies to introduce specific mutations of interest, providing deeper insights into gene functions and interactions within an African genetic context. This unique iPSC line has the potential for broad research applications and will significantly help to bridge the gap in our understanding of the interplay between genetics and disease in underrepresented populations.

## 2. Resource Details

Due to cultural and religious norms surrounding organ donation in most African communities, access to human tissue samples (e.g., post-mortem brain) is limited [5]. Given this challenge, iPSC modelling provides a critical opportunity to produce diverse human cell types in an African genetic background to advance our understanding of diseases. Here, we report the generation of an iPSC line derived from a healthy, 60-year-old male indigenous Nigerian from the Babur ethnic group in Borno state. A forearm skin biopsy was collected from the donor using a Stiefel Biopsy Punch, which was used to generate fibroblasts (Figure 1A). The fibroblasts were expanded and characterised based on the expression of fibroblast markers: Vimentin and Smooth Muscle Actin (Figure 1B). The samples underwent testing for 26 commonly checked pathogens, with results confirming the absence of all these pathogens (Table 1).

**Figure 1:**
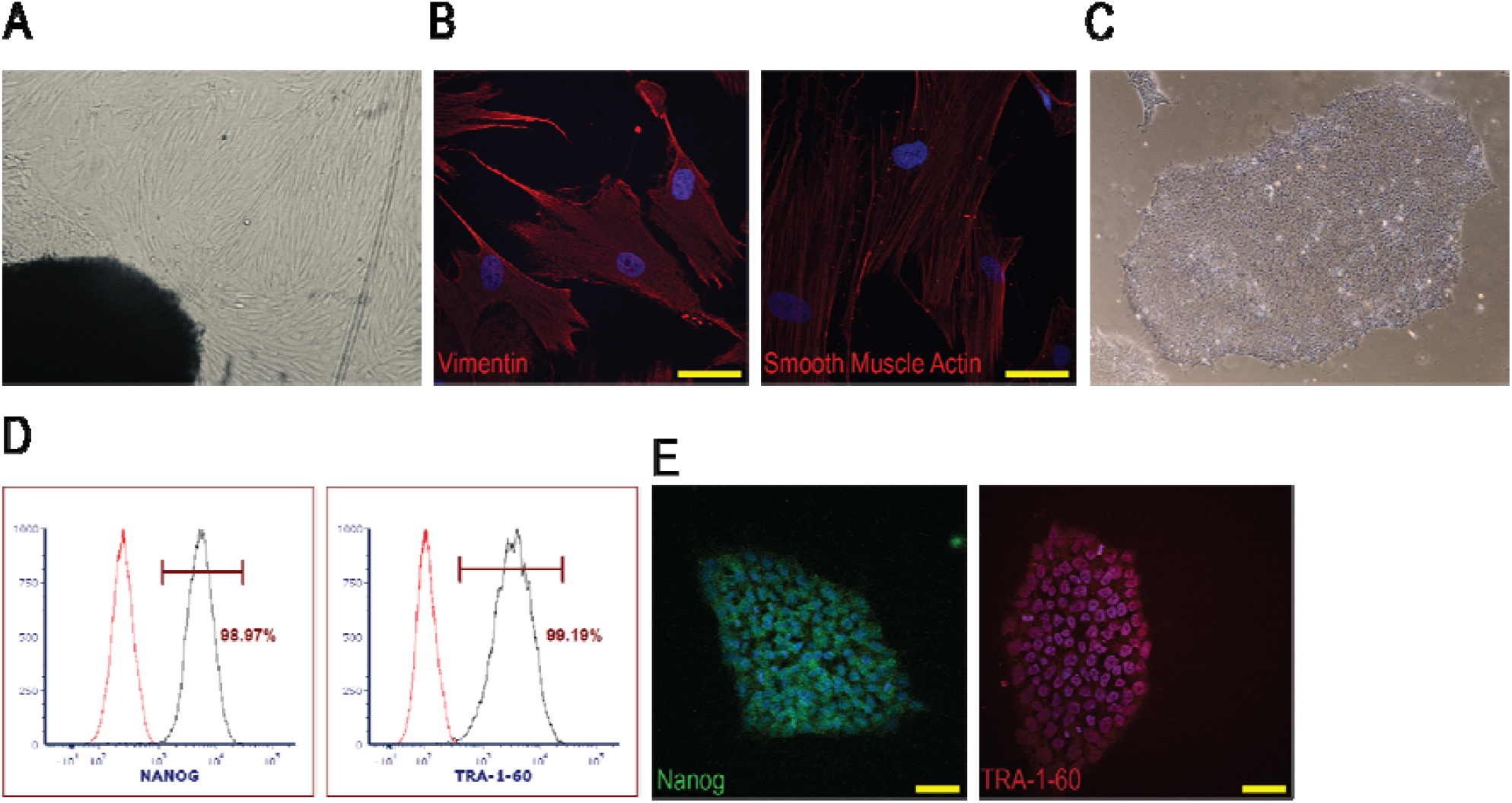
Derivation and Characterization of iPSCs from Fibroblasts. Bright-field image of fibroblasts sprouting from the biopsy (A). Immunofluorescence labelling for Vimentin and Smooth Muscle Actin, markers for fibroblasts (B). Bright-field image of an iPSC, showing a high nuclear-to-cytoplasm ratio and distinct borders (C). Flow cytometry results demonstrating the expression of NANOG and TRA-1-60, markers for iPSCs (D). Immunofluorescence images confirming iPSC identity through labelling for NANOG and TRA-1-60 (E). Scale bar = 50 µm.

**Table 1:** RT-qPCR evaluation of common pathogens in iPSC from healthy Nigerian participant fibroblasts. + = Positive, indicating pathogen presence in sample, - = Negative, indicating pathogen absence from sample

| **Organism of interest** | **Fibroblasts** | **Positive control** | **Negative control** |
| --- | --- | --- | --- |
| Epstein Barr Virus | - | + | - |
| Hantavirus | - | + | - |
| Hepatitis A | - | + | - |
| Hepatitis virus B | - | + | - |
| Hepatitis virus C | - | + | - |
| Herpes Simplex 1 | - | + | - |
| Herpes Simplex 2 | - | + | - |
| HIV-1 | - | + | - |
| HIV-2 | - | + | - |
| Human Adenovirus A | - | + | - |
| Human Adenovirus B | - | + | - |
| Human Adenovirus C | - | + | - |
| Human Adenovirus D | - | + | - |
| Human Adenovirus E | - | + | - |
| Human Adenovirus F | - | + | - |
| Human and Mouse LCMV | - | + | - |
| Human Cytomegalovirus | - | + | - |
| Human Herpesvirus 6 | - | + | - |
| Human Herpesvirus 8 | - | + | - |
| Human Papillomavirus 16 | - | + | - |
| Human Papillomavirus 18 | - | + | - |
| Human T-lymphotropic virus 1 | - | + | - |
| Human T-lymphotropic virus 2 | - | + | - |
| Seoul hantavirus | - | + | - |
| Sin Nombre | - | + | - |
| Varicella | - | + | - |

iPSC reprogramming was achieved using Sendai virus-based delivery of reprogramming factors. Colonies were manually picked, leading to the generation of two stable iPSC clones: OSTiFGM2-A and OSTiFGM2-B. Both clones formed characteristic iPSC morphology with compact colonies with distinct borders and well-defined edges (Figure 1C). Cells from both lines maintained a high nuclear-to-cytoplasm ratio, with a normal growth rate and a subculture frequency of 4-6 days. Subsequently, the iPSC clones were characterised, showing expression of pluripotency markers TRA-1-60 and NANOG, as evidenced by flow cytometry and immunocytochemistry (Fig 1D and E) (Fig 1D and E).

Both the parent fibroblasts and the two iPSC clones were evaluated for chromosomal abnormalities. Copy number variation (CNV) analyses were conducted on DNA isolated from the cells using the Illumina Infinium Global Screening Array. No new CNVs were detected in the two iPSC clones compared to the parental fibroblasts (Figure 2).

**Figure 2:**
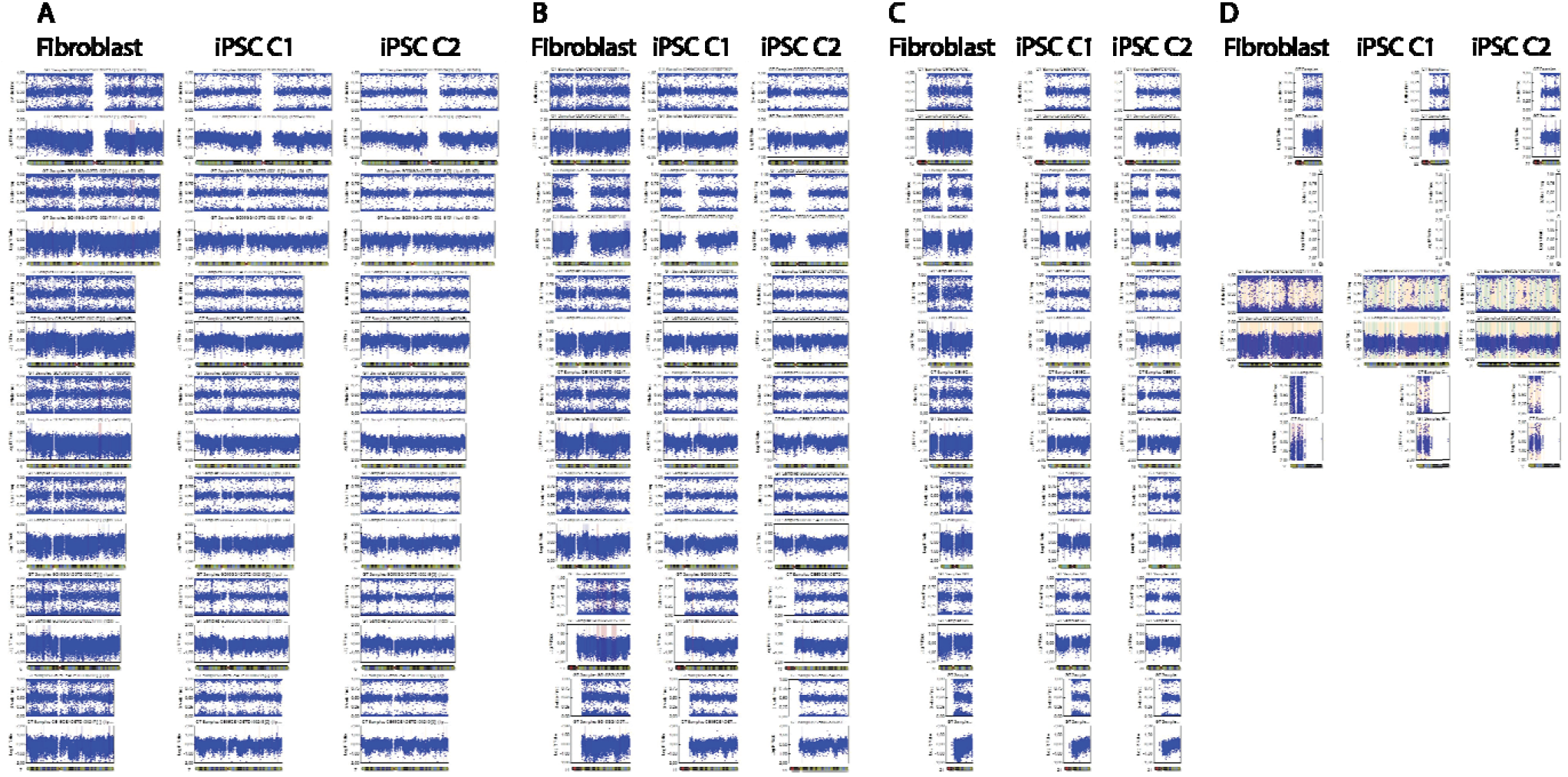
Chromosomal Stability Assessment between Parental Fibroblasts and Derived iPSC Clones. Both B-allele frequency (indicating allele-specific copy number) and log R ratio (indicating total copy number) plots for all chromosomes are presented for the parental fibroblast sample, and iPSC clones A (iPSC C1) & B (iPSC C2). Detailed chromosomal analyses are categorised into chromosomes 1 – 7 (A), 8 – 14 (B), 15 – 21 (C), and 22, X, and Y (D). No major chromosomal abnormalities were identified in iPSC clones when compared to their parental fibroblasts.

To investigate the differentiation potential of the iPSCs, both iPSC clones were first induced into neural progenitor cells (NPCs), as confirmed by PAX6 and Nestin immunoreactivity (Figure 3A). These NPCs were then differentiated further into astrocytes, as evidenced by GFAP staining (Figure 3B). The GFAP labelling showed that the induced astrocytes exhibit varied morphologies, typical of astrocytes.

**Figure 3:**
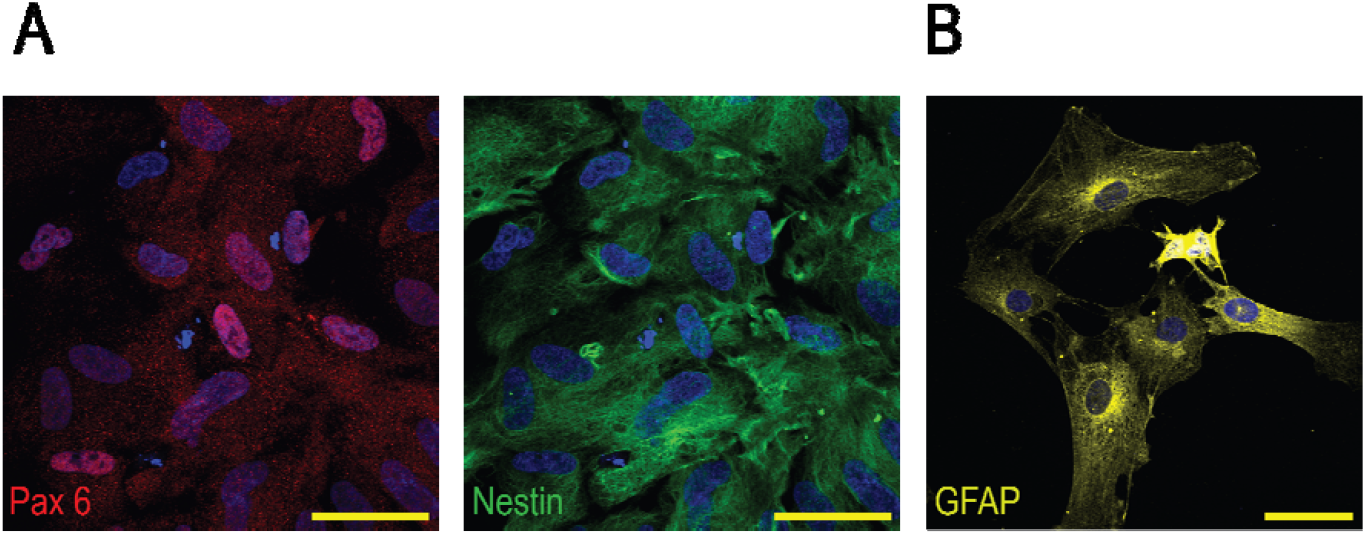
Immunofluorescence labelling showing PAX6 and Nestin, markers of neural progenitor cells (A) and GFAP, a marker of astrocytes (B). Scale bar = 50 µm

## 3. Material and Methods

### 3.1. Biopsy collection and fibroblast generation

A forearm skin biopsy was collected from donors at the University of Maiduguri Teaching Hospital by a surgeon using a 6 mm Stiefel Biopsy Punch, after the area had been anaesthetised. The tissue samples were immediately rinsed twice into two sterile 50 mL falcon tubes containing phosphate-buffered saline (PBS). The biopsies were then placed into Dulbecco’s Modified Eagle Medium (DMEM) supplemented with 1% (v/v) L-glutamate (L-Glu), 1% (v/v) penicillin/streptomycin (Pen/Strep), 2.5% (v/v) HEPES solution, and 10% (v/v) Fetal Calf Serum. The use of HEPES buffer was chosen as a necessary modification to buffer the media due to CO_2_ leakage in the Marte’s Laboratory incubator which had previously led to the loss of biopsy samples. The biopsies were transported for processing at the Marte’s Laboratory at the University of Maiduguri. In the tissue culture fume hood, sterile forceps were used to hold the tissue inside a 6-well plate containing a thin layer of DMEM media to prevent the sample from drying out. A sterile scalpel was used to cut the tissues into 1-2 mm pieces. About 6 - 8 pieces of the cut samples were transferred to a T24 flask without media, with the epidermis side of the tissue facing up, and incubated for 10 - 15 minutes at 37°C to allow tissue samples to adhere to the flask. The flask was then returned to the hood and supplemented with a small amount of media to form a thin layer. The T25 flask lid was tightly closed to prevent air exchange with the external environment and incubated inside the 37°C incubator without CO_2_. Every two days, 50% of the media was gently replaced with fresh media, ensuring that the attached biopsies were not disturbed. Around the sixth day, fibroblasts started sprouting from underneath the biopsy pieces. The cells were expanded until passage six, when enough cells were obtained. To freeze the cells, they were resuspended in the media supplemented with 10% dimethyl sulfoxide (DMSO), transferred into a cryovial, and stored at -20°C due to the unavailability of a -80°C freezer in Marte’s Laboratory. The next day, the vials were transferred into liquid nitrogen storage. Following this, samples were shipped in a dry shipper to the UK for onward processing into iPSCs. The testing of the fibroblasts for the 26 common pathogens was outsourced to Oxford StemTech (Table 1).

### 3.2. iPSC Reprogramming

Initially, we used the ReproRNA-OKSGM cocktail and protocol (STEMCELL Technologies) to reprogram the fibroblasts into iPSCs. Despite the successful reprogramming of a human neonatal foreskin or adult skin (normal) fibroblast line purchased from Sigma (Cat 106-05A), multiple attempts to reprogram the Nigerian fibroblasts failed for reasons that are not entirely clear, prompting the switch to Sendai virus-based ReproPlexTM technology (Oxford StemTech). This was used to reprogram the cells into two iPSC clones (OSTiFGM2-A and OSTiFGM2-B).

#### 3.2.1 iPSC Maintenance and Characterisation

The morphology of the iPSCs was visualised using a brightfield microscope. The iPSCs were maintained in a 37°C / 5% CO_2_ incubator and expanded on Corning® Matrigel® hESC-qualified Matrix (Corning 354277)-coated plates with mTeSR™ Plus medium (STEMCELL Technologies) changes every other day. Upon reaching 80-90% confluency, ReLeSR (STEMCELL Technologies) was used to detach the colonies. The detached colonies were then pelleted at 300g_(av)_ and reseeded with mTeSR Plus medium supplemented with 10 µM Rock Inhibitor (Y-27632, STEMCELL Technologies). At passage 10, RT-qPCR demonstrated effective clearance of the Sendai virus using both positive and negative controls. Free of transgenes, the iPSCs were characterised through flow cytometry and immunofluorescence labelling for the expression of iPSC markers (TRA-1-60 and NANOG).

### 3.3 Mycoplasma test

Media supernatants were harvested from all iPSC lines and assessed for the presence of mycoplasma using Lonza’s MycoAlert kit. The mycoplasma assay confirmed the absence of mycoplasma in all tested lines.

### 3.4 Differentiation to Neural Progenitor Cells

First, 6-well tissue culture-treated plates were coated with Matrigel for 1 hour. iPSCs at about 70-90% confluency were collected using ReLeSR, mixed with DMEM (without supplements), spun at 300g_(av)_ for 3 minutes, and then induced to NPCs using STEMdiff™ SMADi Neural Induction Kit (STEMCELL Technologies). Specifically, the iPSCs were resuspended into single cells in STEMdiff™ Neural Induction Medium + SMADi media supplemented with 10 µM Rock Inhibitor. The cells were seeded at a density of 2 × 10^5^ cells/cm^2^ and incubated at 37°C / 5% CO_2_ with daily media change excluding Rock Inhibitor. After 6 days, when cells reached 70-80% confluency, they were passaged using ACCUTASE™ (STEMCELL Technologies). To do this, the cells were incubated with ACCUTASE for 5 minutes at 37°C, then resuspended in STEMdiff™ Neural Induction Medium + SMADi in a falcon tube and spun at 300g_(av)_ for 5 minutes. The cell pellet was resuspended in the STEMdiff™ Neural Induction Medium + SMADi containing Rock Inhibitor and re-seeded on matrigel-coated plates. After two further passages, NPCs were obtained and expanded in STEMdiff™ Neural Progenitor Medium. Some NPCs were frozen, characterised, or used for differentiation to astrocytes.

### 3.5 Differentiation to Astrocytes

NPCs were seeded onto a 6-well matrigel-coated plate in STEMdiff™ Neural Progenitor Medium at a 1.5 × 105 cells/cm^2^ density. The next day, the media was replaced with STEMdiff™ Astrocyte Differentiation Medium (STEMCELL Technologies). Media change was performed daily. At day six, when the astrocyte precursors reached 80-90% confluence, they were passaged using ACCUTASE and re-seeded on matrigel-coated plates using STEMdiff™ Astrocyte Differentiation Medium. After 2 passages, the astrocyte precursors were seeded on matrigel-coated plates in Astrocyte Maturation Medium (STEMCELL Technologies) at a density of 1.5 × 10^5^ cells/cm^2^. At day 6 days when the astrocytes precursors reach 80-90% confluence, they were passaged using ACCUTASE. The astrocytes were then seeded onto matrigel-coated plates and characterised after 3 days using an antibody against the astrocyte marker – GFAP Monoclonal Antibody.

### 3.6 Immunostaining

Cells were fixed with 4% Paraformaldehyde/PBS for 15min at room temperature (RT), washed three times with PBS and permeabilised for 15min using 0.5% TritionX-100/PBS. Cells were washed thrice with PBS and blocked for 45min using 4% BSA/PBS at RT. They were next incubated in primary antibodies (Table 2) for 45 minutes, then washed three times with PBS and incubated in corresponding secondary antibodies (Table 2) for 45 min. Following three washes, the cells were mounted onto coverslips using ProLong® Gold Antifade Reagent with DAPI (Thermo Fisher Scientific). The slides were left overnight to dry, then imaged using SP8 Leica Laser Scanning Confocal Microscope.

**Table 2:** Antibodies.

|  | **Antibody** | **Dilution** | **Company Cat #** |
| --- | --- | --- | --- |
| **Fibroblast markers** | Rabbit IgG Vimentin | 1:200 | Abcam Cell, Cat. ab254015 |
|  | Smooth muscle Actin | 1:200 |  |
| **Pluripotency markers** | Anti-Hu TRA-1-60 | 1:200 | STEM Cell Technologies, Cat. 60064PE.1 |
|  | Alexa Fluor™ 488 Nanog Monoclonal Antibody | 1:200 | Fisher Scientific, Cat. 15332040 |
| **Neural progenitor marker** | Anti-Human Nestin Antibody | 1:200 | STEM Cell Technologies, Cat. Clone 10C2 |
|  | Recombinant Anti-PAX6 antibody | 1:200 | Abcam, Cat. ab195045 |
| **Astrocyte marker** | Rabbit Polyclonal Antibody to GFAP | 1:200 | EnCor Biotechnology Inc., Cat. RPCA-GFAP |
| **Secondary antibodies** | Goat anti-Mouse IgG (H+L) Highly Cross-Adsorbed Secondary Antibody, Alexa Fluor™ Plus 488 | 1:500 | Thermo Fisher Scientific, Cat. A32723 |
|  | Goat anti-Rabbit IgG (H+L) Cross-Adsorbed Secondary Antibody, Alexa Fluor™ 568 | 1:500 | Thermo Fisher Scientific, Cat. A-11011 |

## Acknowledgement

This work was funded by a Sussex Neuroscience Seed Fund Fellowship, Alzheimer’s Association Research Fellowship, Alzheimer’s Research UK South Coast Network, and Sussex School of Life Sciences Ewart Bequest Fund, to Mahmoud Bukar Maina. We would like to acknowledge the invaluable support and generosity of the donors and staff of the University of Maiduguri Faculty of Pharmacy and Yerwa Express for their efforts in donor recruitment and downstream technical support during the early stages of this work in Nigeria. We are also very grateful to Prof. Marycelin Baba and her staff at the World Health Organization Laboratory at the University of Maiduguri Teaching Hospital for providing us with the technical resources necessary to culture our biopsies in a non-CO2 incubator. iPSC reprogramming with Sendai virus-based ReproPlexTM technology, Sendai virus clearance, mycoplasma testing, and karyotyping analysis were performed by Oxford StemTech.

